# The effect of body mass index on smoking behaviour and nicotine metabolism: a Mendelian randomization study

**DOI:** 10.1101/299834

**Authors:** Amy E. Taylor, Rebecca C. Richmond, Teemu Palviainen, Anu Loukola, Jaakko Kaprio, Caroline Relton, George Davey Smith, Marcus R. Munafò

**Affiliations:** Population Health Sciences, Bristol Medical School, University of Bristol, Bristol, United Kingdom.; NIHR Biomedical Research Centre at the University Hospitals Bristol NHS Foundation Trust and the University of Bristol, United Kingdom.; MRC Integrative Epidemiology Unit at the University of Bristol, Bristol, United Kingdom.; Institute for Molecular Medicine FIMM, Helsinki Institute for Life Science, University of Helsinki, Helsinki, Finland; Department of Public Health, Medical Faculty, University of Helsinki, Helsinki, Finland; UK Centre for Tobacco and Alcohol Studies, School of Experimental Psychology, University of Bristol, Bristol, United Kingdom.

**Keywords:** Body mass index, smoking, ALSPAC, Mendelian Randomisation

## Abstract

**Background:** Given clear evidence that smoking lowers weight, it is possible that individuals with higher body mass index (BMI) smoke in order to lose or maintain their weight.

**Methods and Findings:** We undertook Mendelian randomization analyses using 97 genetic variants associated with BMI. We performed two sample Mendelian randomization analyses of the effects of BMI on smoking behaviour in UK Biobank (N=335,921) and the Tobacco and Genetics consortium genomewide association study (GWAS) (N≤74,035) respectively, and two sample Mendelian randomization analyses of the effects of BMI on cotinine levels (N≤4,548) and nicotine metabolite ratio (N≤1,518) in published GWAS, and smoking-related DNA methylation in the Avon Longitudinal Study of Parents and Children (N≤846).

In inverse variance weighted Mendelian randomization analysis, there was evidence that higher BMI was causally associated with smoking initiation (OR for ever vs never smoking per one SD increase in BMI: 1.19, 95% CI: 1.11 to 1.27) and smoking heaviness (1.45 additional cigarettes smoked per day per SD increase in BMI, 95% CI: 1.03 to 1.86), but little evidence for a causal effect with smoking cessation. Results were broadly similar using pleiotropy robust methods (MR-Egger, median and weighted mode regression). These results were supported by evidence for a causal effect of BMI on DNA methylation at the aryl-hydrocarbon receptor repressor (*AHRR*) locus. There was no strong evidence that BMI was causally associated with cotinine, but suggestive evidence for a causal negative association with the nicotine metabolite ratio.

**Conclusions:** There is a causal bidirectional association between BMI and smoking, but the relationship is likely to be complex due to opposing effects on behaviour and metabolism. It may be useful to consider BMI
and smoking together when designing prevention strategies to minimise the effects of these risk factors on health outcomes.

## Introduction

Smoking and obesity are amongst the leading preventable causes of mortality and morbidity worldwide [1]. Understanding pathways which contribute to these risk factors, and the nature of the relationship between them, is therefore of paramount importance for disease prevention. Observationally, current smoking is often associated with lower body mass index [2]. However, heavy smoking has been found to be associated with higher body mass index (BMI) [2, 3]. Given the clustering of unhealthy behaviours such as smoking, low physical activity and poor diet [4], and the strong links between smoking, obesity and sociodemographic factors [5], establishing the existence of and direction of causality is difficult.

Mendelian randomisation (MR), which uses genetic variants associated with exposures as proxies, can help to overcome problems of confounding and reverse causality because, in theory, genetic variants associated with the exposure of interest should be inherited independently of other genetic variants and environmental factors [6]. There is good evidence from MR studies, using a genetic variant that influences the number of cigarettes consumed per day among smokers, that heavier smoking causes a reduction in body mass index and other measures of adiposity [7–9]. This may be explained by nicotine increasing metabolic rate and/or lowering appetite and therefore changing energy balance [2]. To support this, there is a large body of evidence showing that smoking cessation is accompanied by weight gain [10–15], though with large individual variation in the amount gained.

Given that smoking lowers body weight, it is plausible that the association between BMI and smoking is bidirectional; that is more overweight individuals may take up smoking, smoke more heavily, or continue to smoke rather than quit, in order to lower weight. Weight gain is commonly cited as a concern for smokers who are considering quitting smoking [10]. This has been found most consistently in women [10], although there is also evidence that weight concern is associated with motivation to quit smoking in men [16]. Weight concern or body dissatisfaction amongst adolescents may also increase the likelihood of smoking initiation [17, 18]. However, it is important to note that the relationship between weight concern and BMI is complex; for example, it may be U-shaped in males [19]. Amongst young people in the Avon Longitudinal Study of Parents and Children (ALSPAC), higher BMI was associated with smoking initiation in females, but not in males, whereas body dissatisfaction was associated with higher risk of smoking initiation in both sexes [20]. Smoking and obesity are also both associated with increased risk of anxiety and depression [21] and there is evidence that the link between higher BMI and depressive symptoms is causal [22]. Therefore, it is possible that BMI could lead to smoking through its effects on mental health, although strong evidence of causality between mental health and smoking is yet to be established.

In addition to behavioural links, it is possible that BMI could alter smoking behaviour via physiological effects. Higher BMI could result in lower blood nicotine levels for the same amount smoked, due to higher total blood volume or absorption of nicotine or its metabolites by fatty tissue [23]. It has been demonstrated that BMI is negatively correlated with nicotine levels following administration of nicotine replacement therapy [24]. This could mean that individuals with higher BMI would need to smoke more in order to experience the same effect of nicotine. BMI may also affect nicotine metabolism, which is commonly measured by the nicotine metabolite ratio (NMR). Studies have shown that individuals with higher NMR (reflecting faster metabolism of nicotine) smoke more heavily and are less likely to give up smoking [25, 26]. Observationally, BMI tends to be negatively correlated with NMR [27]. This could plausibly be because NMR lowers BMI through its effect on increasing smoking, although it has been argued that evidence points towards the relationship being in the opposite direction, from BMI to NMR [27].

A previous genetic analysis demonstrated that higher genetically determined BMI was associated with increased likelihood of smoking initiation and higher tobacco consumption [28]. This was interpreted by the authors as shared genetic aetiology for BMI and smoking rather than a causal effect of BMI on smoking. For example, variants in the brain derived neurotrophic factor (*BDNF*) gene associate with both BMI and smoking initiation at genomewide significance level [29, 30]. We sought to extend this work and explore the potential causal effect of BMI on smoking using a larger number of genetic variants and Mendelian randomisation methods which are more robust to potential pleiotropy [31–33]. Using genetic variants associated with BMI from the largest published GWAS of BMI to date [30], we investigated whether BMI causes differences in smoking behaviour and total tobacco exposure, by looking at both self-reported measures of smoking and biological measures of exposure (cotinine and DNA methylation). We also used this approach to investigate whether BMI causally influences NMR. We performed analyses using several datasets: the Tobacco and Genetics Consortium GWAS [29], the Cotinine Consortium GWAS [34] and the largest NMR GWAS conducted to date [35], the UK Biobank [36] and ALSPAC [37].

## Methods

We performed two sample Mendelian randomisation using summary data from GWAS and individual level data from the UK Biobank.

## Study samples

### GWAS summary data: BMI

We obtained summary data on the association of genetic variants with BMI from the most recent GIANT BMI GWAS [30]. We used the 97 independent SNPs identified as reaching genome-wide significance with BMI. Associations between genetic variants and BMI (betas and standard errors) were obtained from the meta-analysis of the European sex-combined datasets (N ≤ 322,135) [30]. A full list of SNPs used in each analysis is shown in Supplementary Table S1.

### GWAS summary data: smoking related outcomes

We obtained estimates (beta coefficients/odds ratios and standard errors) of the association of BMI-related genetic variants with smoking initiation (ever vs never smoking) (N ≤ 74,035), age of initiation (N ≤ 24,114), smoking cessation (former vs current smoking) (N ≤ 41,278) and smoking heaviness amongst ever smokers (cigarettes smoked per day) (N ≤ 38,101) from the Tobacco and Genetics (TAG) consortium GWAS [29]. We looked up associations of BMI-related SNPs with cotinine in summary data from a published GWAS of cotinine levels in current daily cigarette smokers (N ≤ 4,548) [34] and with the NMR in summary data from a GWAS in cotinine-verified current smokers [35]. Summary statistics for the NMR GWAS not adjusted for BMI were obtained from the study authors separately for the Finnish Twin Study (FinnTwin), the Young Finns Study (YFS) and the National FINRISK study.

### GWAS summary data: DNA methylation

We performed genome-wide association analysis of DNA methylation at the aryl-hydrocarbon receptor repressor (*AHRR*) methylation site cg05575921 (the strongest smoking-associated methylation locus identified to date [38]) in the Avon Longitudinal Study of Parents and Children (ALSPAC) ARIES resource [39]. ALSPAC is a longitudinal birth cohort, which recruited 14,541 pregnant women with due dates between 1 April 1991 and 31 December 1992. Information on these women and their children has been collected at clinics and via questionnaires ever since [37, 40]. Please note that the study website contains details of all the data that is available through a fully searchable data dictionary (http://www.bris.ac.uk/alspac/researchers/data-access/data-dictionarv/). Ethical approval for the study was obtained from the ALSPAC Ethics and Law Committee and the Local Research Ethics Committees. The ARIES resource includes 1,018 mother offspring pairs. DNA methylation in the mothers was assessed from blood samples taken at two timepoints: during pregnancy and^∼^18 years later. Genome-wide DNA methylation profiling in ARIES was performed using the Illumina Infinium HumanMethylation450 BeadChip (450ͰK) array [39]. Full details of the GWAS methods are provided in supplementary material. The sample used in the GWAS (N ≤ 846) included smokers and non-smokers. Beta coefficients and standard errors of the association with methylation for each of the BMI-related SNPs were obtained from the GWAS summary statistics.

## UK Biobank

We also used data on individuals from the UK Biobank, which recruited over 500,000 individuals (aged between 40 and 70 years) in the UK [41]. Individuals attended assessment centres between 2006-2010, where they completed a questionnaire on lifestyle factors and had blood samples and measurements taken. Individuals were classified as ever smokers if they had smoked more than 100 cigarettes in their lifetime and current smokers if they indicated that they were still smoking. Cigarettes smoked per day amongst current smokers and past regular smokers was reported on a continuous scale. BMI was calculated as weight(kg)/height(m)^2^. In this analysis, we included unrelated individuals of white British ancestry (N=335,921) (see supplementary material for details).

## Statistical analysis

In two-sample Mendelian Randomisation analysis, we calculated the ratio of the SNP-outcome and SNP-exposure associations (the Wald estimator) for each of the 97 BMI-related SNPs (see Supplementary Table S1), to give an estimate of the effect of BMI on the outcome. Where BMI-related SNPs were not available in the outcome GWAS, proxy SNPs (with an R-squared value of > 0.9 with the original SNP) were used if available. The single SNP estimates were combined in an inverse variance weighted (IVW) random effects meta-analysis, as outlined by Burgess and colleagues [42], using the mrrobust package in Stata [43]. For the analysis of smoking initiation, we excluded the genetic variant in *BDNF*, as this locus is likely to be pleiotropic and is associated with smoking initiation at genome-wide significance level [29].

Within UK Biobank, we generated a weighted BMI genetic risk score from dosage scores of the 97 SNPs, using the weights from the combined ancestries GIANT analysis [30] and tested the association of the standardised risk score against measured BMI using linear regression. We calculated associations of each SNP with smoking behaviour phenotypes using logistic or linear regression, adjusted for 10 principal genetic components, and produced causal estimates using the same two sample MR IVW method as outlined above. We performed primary analyses in the full sample, but also stratified by sex, given evidence from previous literature that the relationship between weight concern and smoking might be stronger in females. Results from TAG and UK Biobank were meta-analysed using inverse variance weighted fixed effects meta-analysis.

We also performed analyses which are more robust to potential pleiotropy, MR Egger [31], weighted median regression [32] and the mode based estimator [33]. The MR Egger method is similar to IVW, but allows the intercept of the regression line to change. The intercept is a test of directional pleiotropy; if the intercept differs from zero, this indicates that there is directional pleiotropy. The slope obtained from MR Egger is an estimate of the causal effect after taking into account this directional pleiotropy [31]. Weighted median regression generates a consistent estimate of a causal effect even when up to 50% of SNPs are invalid instruments [32]. The mode based estimator method assumes that the most commonly occurring causal effect estimate is a consistent estimate of the true causal effect [33].

In addition, we attempted to replicate previous analyses investigating the causal effect of smoking on BMI [7], using the rs16969968 functional variant in the *CHRNA3*-*A5*-*B4* gene cluster, which increases smoking heaviness (cigarettes smoked per day) amongst smokers [44]. We regressed the rs16969968 SNP on BMI in never, former and current smokers, adjusting for age, sex and principal components in UK Biobank.

All analyses were conducted in Stata (version 14.1).

## Results

### Association of BMI genetic risk score with BMI

Within UK Biobank, each SD increase in genetic risk score was associated with a 0.64kg/m^2^ increase in BMI (95% CI: 0.62 to 0.65). There was evidence that the association of the BMI genetic risk differed by smoking status (p for heterogeneity ≤ 0.001), with the strongest association seen in current smokers (Figure S1).

### MR analysis of effect of BMI on self-reported smoking behaviours

There was evidence that BMI was causally associated with increased likelihood of smoking initiation (Figure 1A and Supplementary Tables S1 and S2). In IVW Mendelian randomisation analysis combining the TAG and UK Biobank results, a one SD increase in BMI increased the odds of being an ever rather than a never smoker by 19% (OR: 1.19, 95% CI: 1.11 to 1.27). Findings from weighted median, MR Egger and mode weighted regression were consistent with a positive association with smoking initiation, although magnitudes of association were lower in median and weighted mode regression. In MR Egger analysis, there was no clear evidence for directional pleiotropy.

**Figure 1.**
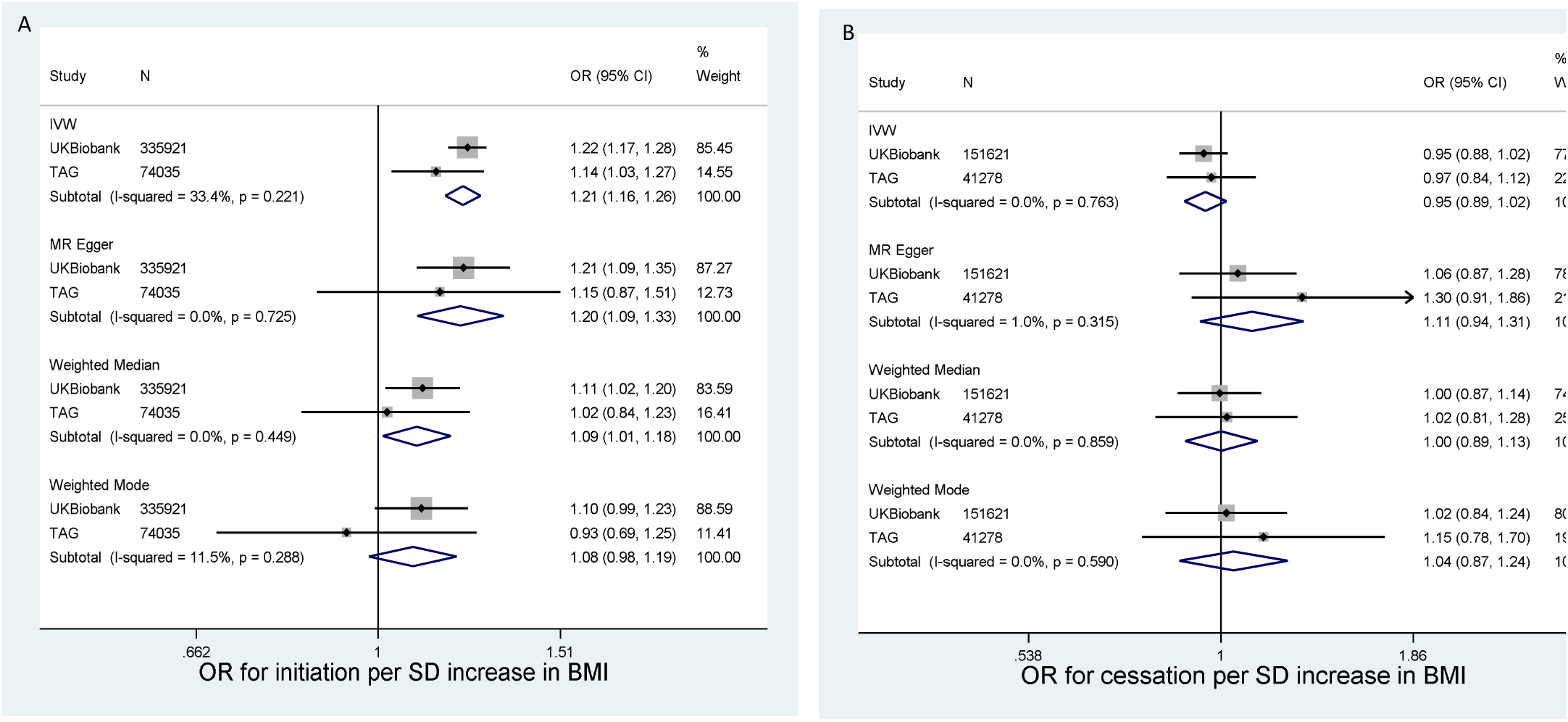
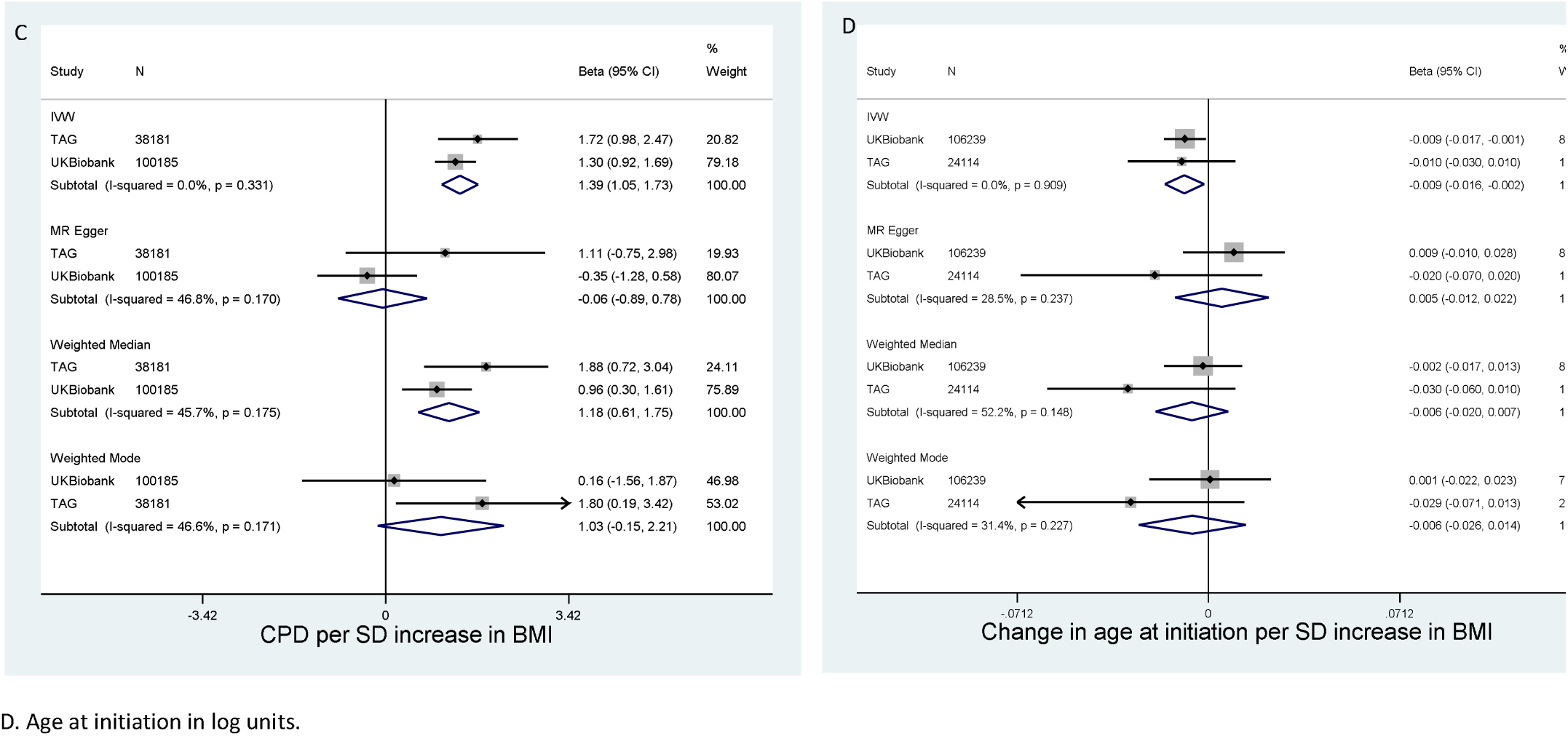
Association between BMI genetic risk score and smoking phenotypes in TAG and UK Biobank.

We also found some evidence for a causal effect of higher BMI on smoking heaviness within smokers (Figure 1C). In IVW analysis, each SD increase in BMI increased smoking heaviness by 1.45 (95% CI: 1.03 to 1.86) additional cigarettes per day. Estimates of these associations were similar for median and weighted mode regression. However, the combined estimate from MR Egger was not consistent with the findings from IVW (β= 0.04, 95% CI −0.94, 1.03).

A one SD increase in BMI was associated with a −0.01 log unit decrease in age at initiation (95% CI: −0.02 to 0.0003) in IVW analysis. Results from the other analytical approaches were consistent with this effect but were imprecise (Figure 1D). There was no clear evidence using any of the approaches for a causal effect of BMI on smoking cessation (Figure 1B).

Results were similar for males and females in UK Biobank (p-values for heterogeneity in comparisons of IVW analyses all >0.6) (Tables S3 and S4).

### MR analysis of effect of BMI on DNA methylation

In the ALSPAC mothers, DNA methylation at *AHRR* was negatively associated with being a smoker and with cigarettes per day (Supplementary Table S5).

There was evidence for a causal effect of BMI on *AHRR* DNA methylation in the ALSPAC mothers in ARIES (Table 1). In IVW Mendelian randomisation analysis, a one SD increase in BMI decreased *AHRR* DNA methylation by 0.33 SD (95% CI: −0.55 to −0.11) in samples taken ^∼^18 years post pregnancy and by 0.23 SD (95% CI: −0.47 to 0.01) in the antenatal samples. Evidence from the pleiotropy robust methods were consistent with the results from IVW analysis, but evidence for associations in the antenatal samples was weak using these approaches.

**Table 1.**
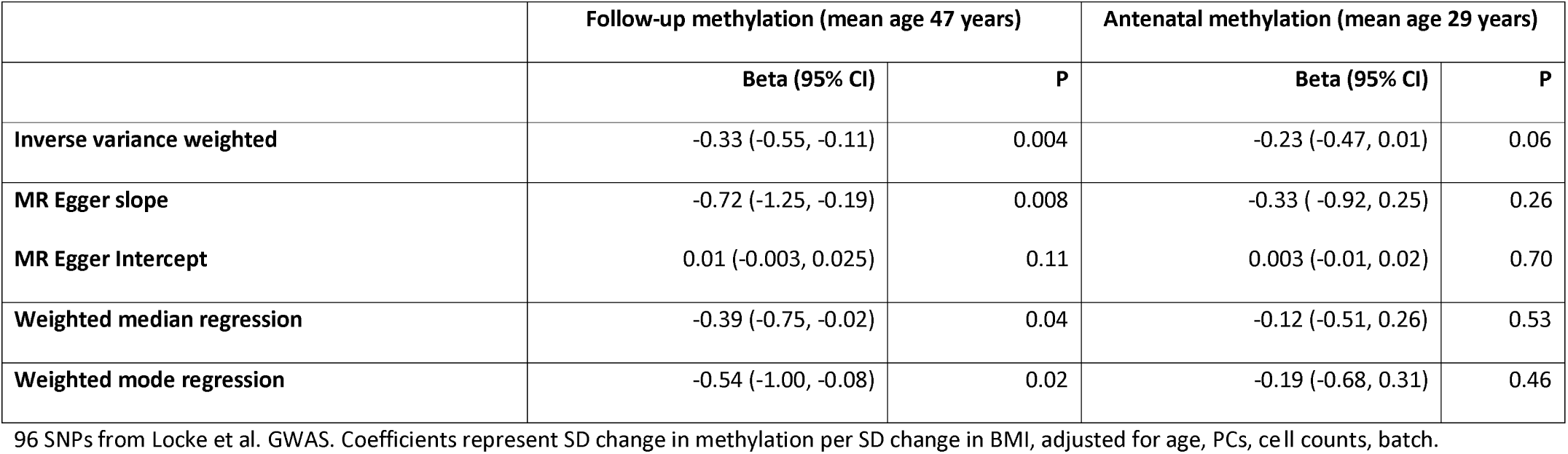
Two sample MR of causal effect of BMI on *AHRR* methylation (cg05575921) in ARIES (N=up to 846)

### MR analysis of effect of BMI on cotinine

Using data from the cotinine GWAS, we found no clear evidence for a causal effect of BMI on cotinine levels (beta from IVW: 0.05 SD, 95% CI: −0.13 to 0.23) (Table 2).

**Table 2.** Two sample MR of causal effect of BMI on cotinine (N=up to 4,548)

|  | Cotinine (SD) |  |
| --- | --- | --- |
|  | Beta (95% CI) | P |
| Inverse variance weighted | 0.05 (-0.13, 0.23) | 0.62 |
| MR Egger slope | 0.02 (-0.41, 0.46) | 0.91 |
| MR Egger Intercept | 0.001 (-0.01, 0.01) | 0.92 |
| Weighted median regression | 0.03 (-0.26, 0.32) | 0.84 |
| Weighted mode regression | -0.005 (-0.370, 0.360) | 0.98 |

Using 95 SNPs from Locke et al. GWAS. Coefficients represents SD change in cotinine per SD change in BMI. I-squared for heterogeneity in IVW analysis:
0%, p-value=0.87

### MR analysis of effect of BMI on nicotine metabolite ratio

Across the FinnTwin, FINRISK and YFS studies, there was suggestive evidence that higher BMI was associated with lower NMR (-0.47 per SD increased in BMI, 95% CI: −0.78, −0.12 in IVW analysis). The magnitude of association was consistent across the other approaches; however, there was a large amount of heterogeneity between the studies for weighted median and weighted mode analyses. Clear evidence for a negative association between BMI and NMR was only seen in the FinnTwin study (Supplementary Table S6).

**Figure 2.**
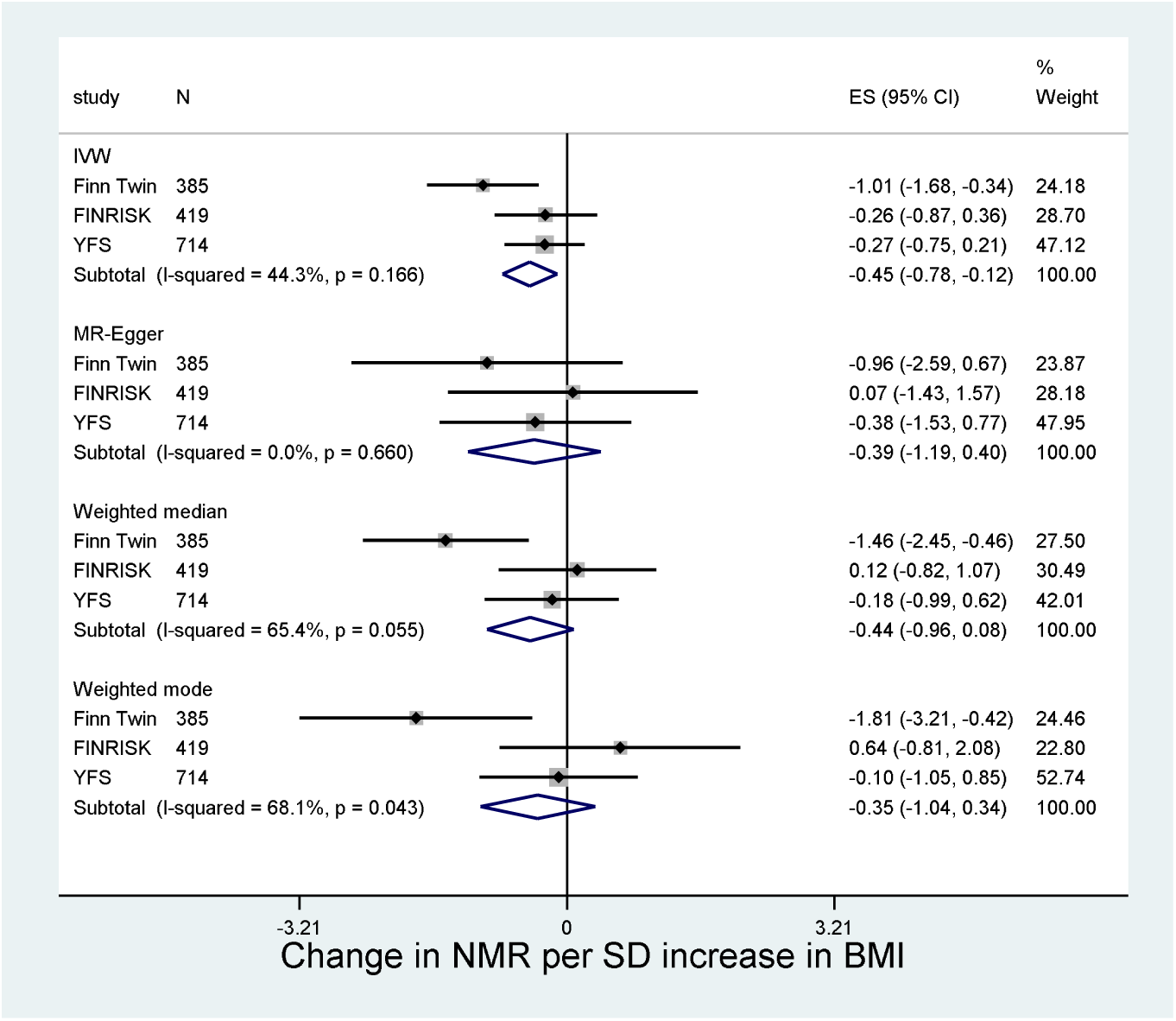
Two sample MR of effect of BMI on NMR in FinnTwin, FINRISK and YFS.

### MR analysis of the effect of smoking heaviness on BMI

Consistent with previous studies [7, 8], the minor allele of the smoking heaviness related variant, rs16969968, increased number of cigarettes smoked per day by 0.95 (95% CI: 0.79 to 1.11, N = 22,568) and decreased BMI in current (beta per minor allele: −0.21, 95% CI: −0.29 to −0.13, N = 32,685), but not former (beta per minor allele: 0.01, 95% CI: −0.03 to 0.05, N = 116,158) or never smokers (beta per minor allele: 0.02, 95% CI: −0.02 to 0.05, N = 181,333, p for interaction between smoking groups<0.001) in UK Biobank.

## Discussion

Using data from multiple cohorts, we found evidence that higher BMI increases the likelihood of becoming a smoker and increases smoking heaviness within current smokers. This finding was supported by the negative association between the BMI genetic risk score and *AHRR* methylation (which is hypomethylated among smokers). However, the BMI genetic risk score was not associated with cotinine levels and showed some evidence of a negative association with the nicotine metabolite ratio, which we might expect to reduce cigarette consumption [25]. In agreement with previous findings [7], we showed that heavier smoking lowers BMI. Taken together, these results suggest that there may be causal bidirectional associations between smoking phenotypes and BMI, and that these may act in opposing directions.

Our results for smoking initiation and cigarettes per day are similar to those presented by Thorgeirsson and colleagues, who used the TAG dataset but only 32 BMI-related genetic variants, from an earlier GWAS [28]. It is possible that, as they suggest, the effects observed here represent a shared genetic aetiology between BMI and smoking behaviour. However, our results for smoking initiation and cigarettes per day were supported by methods which are more robust to the pleiotropy assumption, MR-Egger, and weighted median and weighted mode MR, giving weight to the explanation that this finding represents a causal effect of BMI on smoking uptake and heaviness. This was supported by the negative association we observed between the BMI genetic risk score and DNA methylation at *AHRR*, given that smoking is associated with lower DNA methylation at *AHRR* [38]. Our finding could, in part, explain the positive association found between the BMI genetic risk score and certain types of lung cancer [45]. Although associations via smoking were ruled out in this analysis, sample sizes for testing associations with smoking behaviour were small.

We did not find clear evidence for an effect of BMI on cotinine levels, which might be expected if having higher BMI increases number of cigarettes smoked per day (and therefore total tobacco intake). It is possible that whilst BMI increases total tobacco intake and therefore absolute cotinine levels, individuals with higher BMI have lower blood cotinine concentration due to higher total blood volume (meaning that cotinine is more diluted in the blood) or greater absorption of cotinine by adipose tissue [23]. These opposing effects could lead to a negligible net effect of BMI on cotinine levels.

We observed some evidence for a causal negative effect of BMI on the nicotine metabolite ratio, although findings should be interpreted with caution as they were very heterogeneous between studies. Although this does not rule out an effect of NMR on BMI mediated through higher tobacco intake, our data provide some support for BMI lowering NMR, the direction hypothesised by Chenoweth and colleagues [27]. Given that it is unlikely that BMI affects plasma cotinine and 3Ͱhydroxycotinine differentially, this could point to an effect of BMI on the enzymes which metabolise these compounds or to indirect effects of BMI via other factors which may affect NMR (e.g. alcohol consumption, hormone levels) [27]. Our findings in relation to NMR demonstrate the potential complexity of the BMI-smoking relationship, with opposing effects on behaviour and metabolism. However, an overall positive effect of BMI on tobacco consumption implies that individuals with higher BMI are still at higher risk of increased tobacco consumption (and therefore the harmful effects of tobacco smoke), even if having higher BMI may reduce levels of metabolites.

Although we have attempted to explore both behaviour and metabolism in our analyses, it is not clear what the mechanisms underlying the association between higher BMI and smoking initiation and cigarette consumption are. If this is due to individuals with higher BMI having greater concerns about weight control, we might also expect to observe evidence for a causal effect with smoking cessation as fear of weight gain is often provided as a reason for continuing to smoke [10]. Importantly, interventions which incorporate weight gain concerns or which aim to tackle weight gain at the same time as smoking cessation may still be effective as weight concerns are not always strongly correlated with or may have non-linear relationships with BMI [19]. Given that there is evidence that higher BMI is causally related to lower socioeconomic status, income and educational attainment [46] and that lower educational attainment causes increased smoking [47, 48] it is possible that any effect of BMI on smoking could be via these sociodemographic factors.

There are several limitations to this analysis. Firstly, there is sample overlap between the BMI GWAS and the smoking, cotinine, NMR and GWA studies (estimated to be up to 17%), which may have biased the results of our two sample Mendelian Randomization analyses in the direction of the observational estimates [49]. However, results for smoking behaviour from UK Biobank (which was not included in the BMI GWAS) were highly consistent with those from TAG, suggesting that these results were not driven by bias due to participant overlap. We also repeated the TAG, cotinine and NMR analyses using beta coefficients and standard errors for BMI generated in UK Biobank and these were similar (data not shown). Secondly, we were unable to test associations of the BMI genetic risk score with BMI in the outcome datasets in the two sample MR. We found some evidence that the association of the BMI genetic risk score with BMI is stronger in current than in former or never smokers in UK Biobank. Therefore effect sizes should be treated with some caution.

In conclusion, our findings support of a bidirectional association between BMI and smoking behaviour. Higher BMI leads to increased likelihood of smoking and greater tobacco consumption, but smoking also serves to reduce BMI. Given that BMI and smoking are both major risk factors for disease, this bidirectional causal relationship highlights the need to consider both of these together in prevention strategies. If having higher BMI does increase smoking, interventions aimed at reducing BMI may also help to prevent smoking uptake.

## Acknowledgements

We are extremely grateful to all the families who took part in this study, the midwives for their help in recruiting them, and the whole ALSPAC team, which includes interviewers, computer and laboratory technicians, clerical workers, research scientists, volunteers, managers, receptionists and nurses. The UK Medical Research Council and Wellcome (Grant ref: 102215/2/13/2) and the University of Bristol provide core support for ALSPAC. This publication is the work of the authors and Amy Taylor and Marcus Munafò will serve as guarantors for the contents of this paper. MRM is a member of the UK Centre for Tobacco Control Studies, a UKCRC Public Health Research: Centre of Excellence. Funding from British Heart Foundation, Cancer Research UK, Economic and Social Research Council, Medical Research Council, and the National Institute for Health Research, under the auspices of the UK Clinical Research Collaboration, is gratefully acknowledged. This work was supported by the Medical Research Council (MC_UU_12013/1, MC_UU_12013/2, MC_UU_12013/6),the NIHR Biomedical Centre at the University Hospitals Bristol NHS Foundation Trust and Cancer Research UK (C18281/A19169 and C57854/A22171) The views expressed in this paper are those of the authors and not necessarily the MRC, Wellcome, NIHR or any other funders.

